# Transsynaptic modulation of presynaptic short-term plasticity induction in hippocampal mossy fiber synapses

**DOI:** 10.1101/2020.10.25.353953

**Authors:** David Vandael, Yuji Okamoto, Peter Jonas

**Author notes:** These authors contributed equally (David Vandael, Yuji Okamoto). Corresponding authors. or, **Corresponding author:** Dr. David Vandael, Dr. Peter Jonas, IST Austria (Institute of Science and Technology Austria), Cellular Neuroscience, Am Campus 1, A-3400 Klosterneuburg, Austria.

## Abstract

The hippocampal mossy fiber synapse is a key synapse of the trisynaptic circuit of the hippocampus. Post-tetanic potentiation (PTP) is the most powerful form of plasticity at this synaptic connection. It is widely believed that mossy fiber PTP is an entirely presynaptic phenomenon, implying that PTP induction is input-specific, and requires neither activity of multiple inputs nor stimulation of postsynaptic neurons for induction. Thus, mossy fiber PTP appears to lack cooperativity and associativity that characterize other forms of plasticity. To directly test these predictions, we made paired recordings between single mossy fiber terminals and postsynaptic CA3 pyramidal neurons in rat brain slices. By stimulating parallel but non-overlapping mossy fiber bouton (MFB) inputs converging onto single CA3 neurons, we confirmed that PTP was input-specific and non-cooperative. Unexpectedly, mossy fiber PTP showed anti-associative induction properties. Mossy fiber excitatory postsynaptic currents (EPSCs) showed only minimal PTP after combined pre- and postsynaptic high-frequency stimulation (HFS) with intact postsynaptic Ca^2+^ signaling (0.1 mM EGTA), but marked PTP in the absence of postsynaptic spiking and after suppression of postsynaptic Ca^2+^ signaling (10 mM EGTA). PTP was rescued by blocking Ca^2+^ entry via voltage-gated R-type and to a smaller extent L-type Ca^2+^channels. PTP was also recovered by extracellular application of group II metabotropic glutamate receptor (mGluR) antagonists and vacuolar-type (v-type) H^+^-ATPase inhibitors, suggesting the involvement of retrograde vesicular glutamate signaling. Transsynaptic regulation of PTP induction may increase the computational power of mossy fiber synapses, and implement a break on hippocampal mossy fiber detonation.

## INTRODUCTION

The hippocampal mossy fiber synapse is a key synapse of the trisynaptic circuit of the hippocampus (Brown and Johnston, 1983; Jonas et al., 1993; Salin et al., 1996; Henze et al., 2000; Nicoll and Schmitz, 2005). Post-tetanic potentiation (PTP) is the most powerful form of plasticity at this synaptic connection (Griffith, 1990; Langdon et al., 1995; Salin et al., 1996; Toth et al., 2000; Vyleta et al., 2016). We previously showed that PTP depends on vesicle pool refilling during its induction phase, leading to an increase in the size of the readily releasable pool (Vandael et al., 2020). Moreover, extra vesicles can be stored for extended periods of time that match *in vivo* recorded inter-event intervals (IEIs) of active dentate granule cells (Vandael et al., 2020). Thus, PTP might be an important mechanism in the formation of short-term memory engrams. However, to define the computational power of this storage mechanism, the precise induction rules need to be determined.

It is widely believed that both induction and expression of mossy fiber PTP are entirely presynaptic (Henze et al., 2000; Nicoll and Schmitz, 2005). This would imply that induction is synapse-specific, and requires neither cooperative activation of multiple inputs nor associative stimulation of postsynaptic neurons (Zalutsky and Nicoll, 1992). However, direct experimental mapping the precise induction rules of mossy fiber PTP has been difficult, because single inputs cannot be reliably stimulated. To measure PTP induction rules at hippocampal mossy fiber synapses, we performed paired recordings between hippocampal mossy fiber terminals and postsynaptic CA3 pyramidal neurons. We found that mossy fiber PTP showed synapse specificity and lack of cooperativity, as predicted (Henze et al., 2000; Nicoll and Schmitz, 2005). However, PTP not only lacked associativity (Schmitz et al., 2003; Sastry et al., 1986; Erickson et al., 2010; Fiebig and Lanser, 2017), but rather showed anti-associative induction properties. Postsynaptically shaped induction rules may help to prevent excessive detonation, increasing the computational power of mossy fiber synapses in the hippocampal network.

## RESULTS

### Specificity and non-cooperativity of mossy fiber PTP

Synapse specificity, cooperativity, and associativity are hallmark properties of induction of synaptic plasticity at glutamatergic synapses. However, rigorous testing of these properties is often difficult, requiring defined stimulation of individual synaptic inputs. To test specificity and cooperativity of PTP at the hippocampal mossy fiber synapse, we combined tight-seal cell-attached stimulation of single mossy fiber terminals (Vyleta et al., 2016; Vandael et al., 2020), with interleaved extracellular stimulation of the mossy fiber tract (Fig. 1a–c) in rat brain slices. To measure excitatory postsynaptic currents (EPSCs) in CA3 pyramidal neurons, postsynaptic neurons were held under voltage-clamp conditions at −70 or −80 mV throughout the entire recording (Fig. 1a–d).

**Fig. 1.**
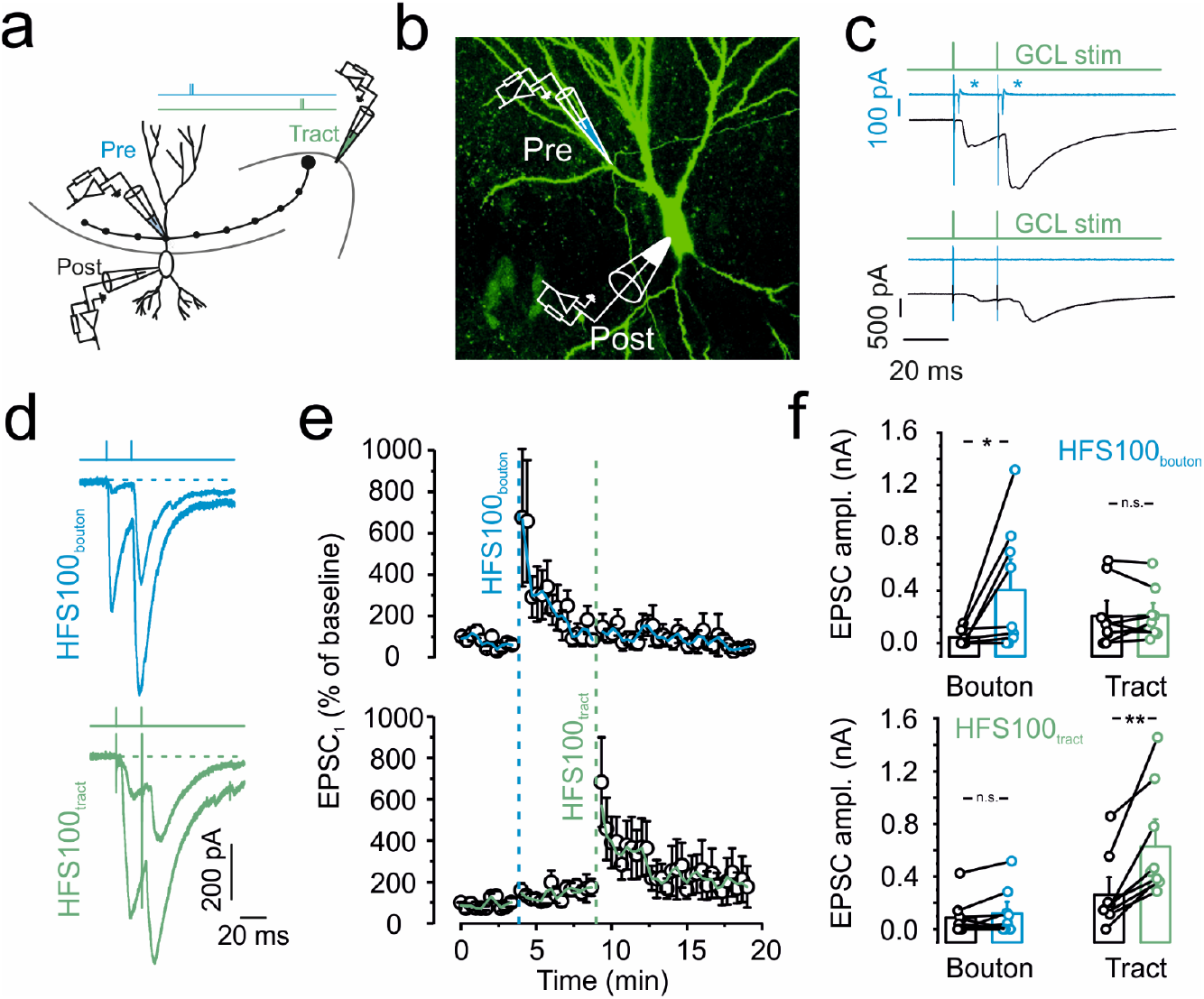
Specificity and non-cooperativity of hippocampal mossy fiber PTP. **a,** Schematic illustration of experimental setup. EPSCs were evoked by alternating stimulation of presynaptic terminals in the minimally invasive tight-seal bouton-attached configuration (blue pipette on proximal apical dendrite) or by tract stimulation (green pipette near granule cell layer). **b,** Maximum stack projection of confocal images of a biocytin-filled pair, stained with Alexa 488-conjugated streptavidin. Pipette on top indicates the approximate location of the tight-seal cell-attached MFB recording, the pipette at the bottom shows wholecell CA3 pyramidal cell recording. **c,** Selecting parallel and independent mossy fiber inputs converging on a single CA3 pyramidal cell. Top, example in which presynaptic action current analysis indicates input overlap. Green trace indicates tract stimulation, blue trace represent currents measured in tight-seal configuration at the level of the MFB, and black trace shows EPSCs. The occurrence of action currents at the MFB (indicated by blue asterisks, top panel) indicates stimulation overlap and led to the exclusion of the recordings. Bottom, example in which presynaptic action current analysis indicates input separation. **d,** Representative traces showing EPSCs evoked by bouton stimulation in tight-seal cell-attached mode (top, blue traces) or by mossy fiber tract stimulation (bottom, green traces). **e,** Normalized EPSC_1_ amplitude plotted against experimental time. Blue vertical dashed line indicates delivery of a 1 s HFS_100_ at the level of the MFB, held in tight-seal cell-attached mode. Green vertical dashed line indicates delivery of HFS_100_ at the level of the mossy fiber tract. Top graph shows unitary EPSCs evoked by direct MFB stimulation, blue line shows running average from 2 points. Bottom graph shows compound EPSCs evoked by stimulation of the mossy fiber tract, green line shows running average. **f,** Summary bar graphs showing the EPSC amplitudes derived from direct bouton stimulation (left, “Bouton”) and from mossy fiber tract stimulation (right, “Tract”). Effect of bouton (top panel) or tract (bottom panel) HFS_100_ as indicated. Boxes represent mean ± SEM, a paired non-parametric Wilcoxon signed rank test was used to test for statistical significance. * indicates P < 0.05, ** represents P < 0.01, and n.s. denotes non-significant difference (P ≥ 0.05).

To test cooperativity, we compared the magnitude of PTP between unitary EPSCs evoked by single-terminal stimulation and compound EPSCs evoked by axon tract stimulation. HFS_100_ applied to a single mossy fiber terminal induced robust PTP of mossy fiber terminal-evoked unitary EPSCs (43.8 ± 22.5 pA in control conditions and 404.5 ± 177.4 pA 20 s after HFS_100_; 9 pairs; P = 0.0179; Fig. 1d–f). Similarly, HFS_100_ applied to the mossy fiber tract induced reliable PTP of tract stimulation-evoked compound EPSCs (260.7 ± 95.4 pA in control conditions and 628.4 ± 146.3 pA 20 s after HFS_100_; P = 0.007; Fig. 1d–f). EPSCs evoked by axon tract stimulation under control conditions had, on average, a 4.7-fold larger amplitude than EPSCs evoked by presynaptic terminal stimulation, verifying the stimulation of multiple synaptic inputs.

To test for synapse specificity, we examined the effects of HFS_100_ applied at the mossy fiber tract on EPSCs evoked by single-terminal stimulation. Independence of the two inputs was verified by continuous monitoring of action currents at the presynaptic terminal (Fig. 1c; 2 out of 11 recordings had to be excluded based on this criterion). HFS_100_ applied to the mossy fiber tract resulted in an only minimal change of the amplitude of presynaptic terminal-evoked EPSCs (86.7 ± 51.7 pA in control and 119.6 ± 66.6 pA 20 s after HFS_100_; 9 pairs; P = 0.176; Fig. 1d–f). Similarly, HFS_100_ applied to the presynaptic terminal left the amplitude of the mossy fiber tract-evoked EPSCs unchanged (206.1 ± 88.2 pA in control and 211.3 ± 66.2 pA 20 s after HFS_100_; 9 pairs; P = 0.678; Fig. 1d–f). Taken together, these results demonstrate that PTP is a synapse-autonomous and synapse-specific phenomenon.

### Anti-associativity of mossy fiber PTP

We next asked whether PTP induction was associative or not (Sastry et al., 1986; Erickson et al., 2010; see Schmitz et al., 2003; Fig. 2). To test the effects of postsynaptic spiking, the postsynaptic CA3 pyramidal neuron was held in the current-clamp configuration during induction (Fig. 2). Since a rise in postsynaptic Ca^2+^ might be required, we used a postsynaptic internal solution containing 0.1 mM EGTA. Surprisingly, PTP evoked by presynaptic HFS_100_ paired with postsynaptic HFS_100_ in current-clamp mode, was absent (98.9 ± 17.7%; 10 pairs; Fig. 2a–c). These results not only indicate that postsynaptic action potentials (APs) are not required for PTP induction, but further suggest that postsynaptic activity suppresses PTP induction, implying an anti-associative induction mechanism (Fig. 2a–c).

**Fig. 2.**
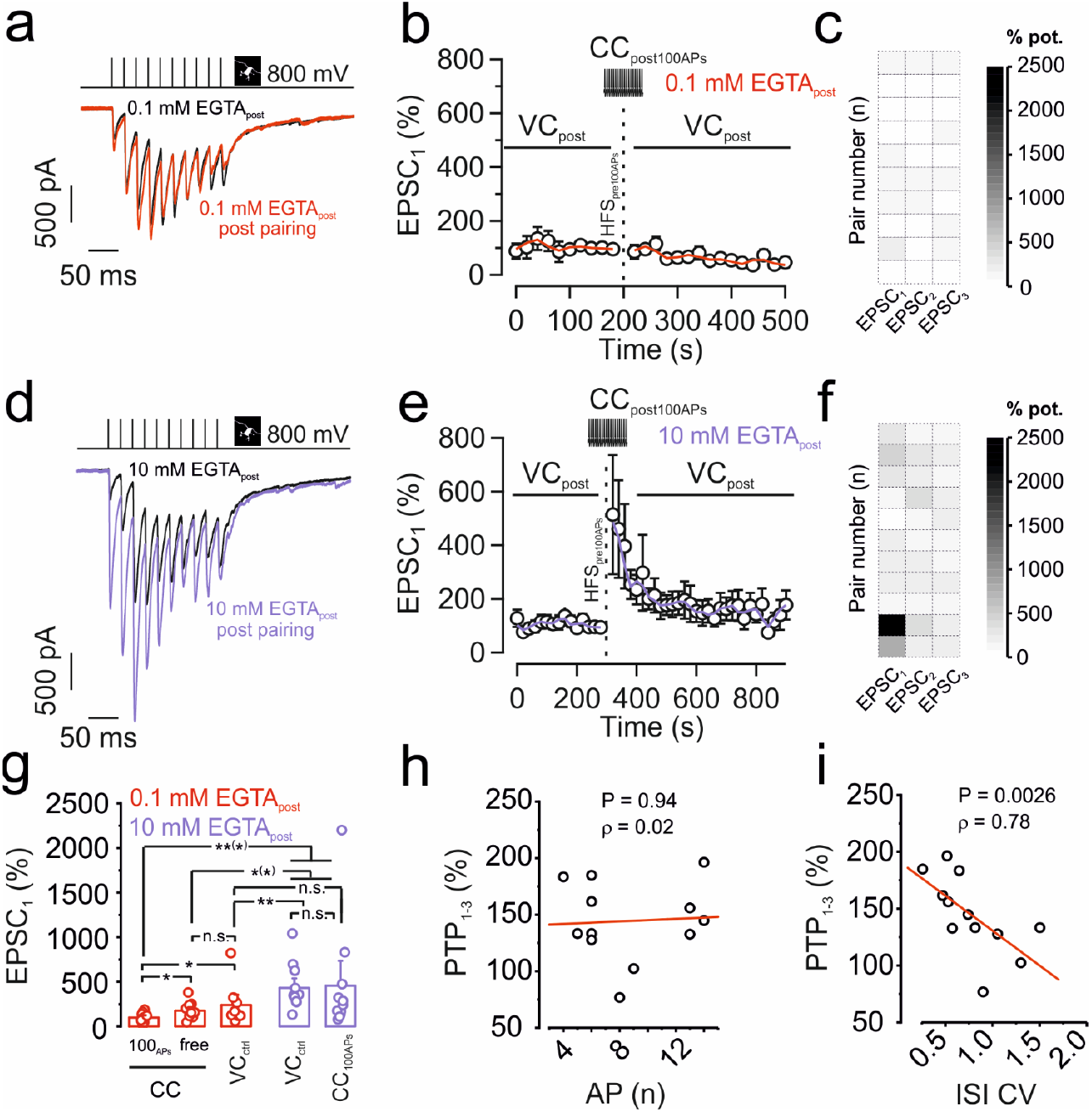
Anti-associative induction properties of mossy fiber PTP. **a–c**, Pre + post HFS_100_ fails to induce PTP with 0.1 mM EGTA in the postsynaptic pipette. **a,** Representative traces of control (black) and post HFS_100_ pairing (red). **b,** Normalized EPSC_1_ amplitude plotted versus experimental time. Black vertical dashed line indicates delivery of an HFS_100_ stimulation (1 s, 100 Hz) in tight-seal cell-attached mode at the level of the MFB, combined with pairing 100 postsynaptic APs in currentclamp mode. Red line shows the running average. **c,**Intensity map showing the degree of potentiation of the first three EPSCs (in %, with control values plotted as 100%) of each single pair between an MFB and a CA3 pyramidal cell, upon pairing with 0.1 mM EGTA postsynaptically. **d–f,** Pre + post HFS_100_ triggers full PTP with 10 mM EGTA in the postsynaptic pipette. **d,** Representative traces of control (black) and post HFS_100_ pairing (red). **e,** Normalized EPSC_1_ amplitude plotted versus experimental time. Black vertical dashed line indicates delivery of an HFS_100_ stimulation (1 s, 100 Hz) in tight-seal cell-attached mode at the level of the MFB, combined with pairing 100 postsynaptic APs in currentclamp mode. Violet line shows the running average. **f,** Intensity map showing the degree of potentiation of the first three EPSCs (in %) of each MFB–CA3 pyramidal neuron pair, with 10 mM EGTA postsynaptically. **g,** Summary bar graphs showing the % of potentiation of EPSC_1_ by pairing 100 presynaptic APs with or without postsynaptic APs with different concentrations of the Ca^2+^ chelator EGTA. Red bar graphs represent pairs where 0.1 mM EGTA was included in the postsynaptic pipette. Violet bar graphs represent pairs where 10 mM EGTA was included in the postsynaptic pipette. CC_100APs_ refers to a–c and d–f (above). CC_free_ refers to experiments where the postsynaptic cell was allowed to spike freely in current-clamp mode upon presynaptic HFS_100_. VC_ctrl_ in the presence of 10 mM EGTA refers to data published in Vandael et al., 2020, where the postsynaptic cell was held in voltage-clamp mode to prevent spiking. Boxes represent mean ± SEM, a non-paired non-parametric Mann-Whitney U test was used to test for statistical significance. * indicates P < 0.05, ** P < 0.01, *** P < 0.001, and n.s. denotes non-significant difference (P ≥ 0.05). **h,** Scatter plot of magnitude of PTP against number of APs of the postsynaptic CA3 pyramidal neuron. The degree of PTP refers to the average PTP obtained for the first three EPSCs (EPSC_1-3_). Data points taken from postsynaptic CCfree recordings upon presynaptic HFS_100_. Note the lack of correlation. Line represents linear regression to the data points (Spearman rank correlation coefficient ρ = 0.02; P = 0.94; 12 pairs). **i,** Scatter plot of magnitude of PTP against coefficient of variation (CV) of ISI duration of the postsynaptic CA3 pyramidal neuron. The degree of PTP refers to the average PTP obtained for the first three EPSCs (EPSC_1-3_). Data points taken from postsynaptic CCfree recordings upon presynaptic HFS_100_. Note a significant negative correlation. Line represents linear regression to the data points (Spearman rank correlation coefficient ρ = 0.78; P = 0.0026; 12 pairs).

To test the possible role of a rise in postsynaptic Ca^2+^, we repeated the associativity experiment (presynaptic HFS_100_ paired with postsynaptic HFS_100_) with a postsynaptic intracellular solution containing 10 mM of the Ca^2+^ chelator EGTA. In the presence of 10 mM postsynaptic EGTA, pairing led to a strong increase of the EPSC_1_ amplitude by 456.9 ± 195.1 % (11 pairs; Fig. 2d–f). The magnitude of PTP was comparable (P = 0.08) to the condition where postsynaptic APs were omitted by voltage clamping of the postsynaptic CA3 cell with a 10 mM EGTA-containing intracellular solution (Fig. 2g). These results demonstrate that postsynaptic activity of the CA3 pyramidal neuron apparently triggers an inflow of Ca^2+^, putting a break on PTP.

To further test the associativity of PTP with a more physiological induction paradigm, we performed a series of experiments in which we allowed the postsynaptic CA3 cell to spike freely (Fig. 2g). The number of postsynaptic APs observed after presynaptic HFS_100_ was 8.7 ± 1.5 on average (ranging between 4 and 14), and also in this setting PTP (177.2 ± 27.02%; 12 pairs) was significantly reduced as compared to control experiments performed in previous work (432.5 ± 73.9%; 12 pairs; P = 0.0005; Fig. 2g; Vandael et al., 2020). However, the magnitude of PTP was larger than in the presynaptic HFS_100_ + postsynaptic HFS_100_ experiments (0.1 mM EGTA post; P = 0.016; Fig. 1g). Thus, few postsynaptic spikes, not locked in time to presynaptic spikes, are sufficient to inhibit PTP, although the degree of inhibition was smaller than with 100 postsynaptic APs. To further characterize the effects of postsynaptic firing on PTP, we plotted the magnitude of PTP against the number of APs and the coefficient of variation of interspike intervals (ISIs). Given that PTP in control conditions affects the first 3 EPSCs (Fig. 2c,f), we used the average PTP value (PTP_1-3_) for quantification. Spearman’s rank correlation analysis revealed that the amount of PTP_1-3_ was not correlated with AP number (ρ = 0.02; P = 0.94), but significantly correlated with the coefficient of variation (CV) of the ISI (ρ = 0.78; P = 0.0026). Thus, patterned forms of postsynaptic activity (i.e. burst firing) may boost postsynaptic Ca^2+^ entry that can trigger anti-associativity of PTP. Given the lack of correlation between the number of post-synaptic spikes and the block of PTP, we subsequently tested whether postsynaptic APs are needed at all to describe the above mentioned phenomenon. We therefore loaded the postsynaptic CA3 pyramidal cell with 0.1 mM EGTA, kept the neuron in voltage-clamp mode, and tried to trigger PTP with our standard induction protocol of 100 APs delivered presynaptically at 100 Hz (no pairing). PTP induced in voltage-clamp conditions with 0.1 mM EGTA was significantly larger (239.5 ± 82.1%; 9 pairs) than PTP with 100 APs in current clamp (P = 0.022; Fig. 2g), smaller than PTP with voltage-clamp induction and comparable to PTP in current clamp with 100 APs with 10 mM EGTA postsynaptically (P = 0.006 and P = 0.145 respectively, Fig. 2g). Taken together, our results suggest that patterned forms of subthreshold postsynaptic activity, independent of the number of Na^+^ spikes, are sufficient to put a brake on mossy fiber PTP.

### Mechanisms underlying anti-associative PTP

To elucidate the signaling mechanisms underlying anti-associative PTP induction, we tested the involvement of pre- and postsynaptic signaling cascades. We first tried to determine the source for the rise in postsynaptic Ca^2+^ (Fig. 3). As N-methyl-D-aspartate (NMDA)-type glutamate receptors are coincidence detectors, and are also present at mossy fiber–CA3 synapses (Jonas et al., 1993; Weisskopf and Nicoll, 1995; Kwon and Castillo, 2008), we first tested the effects of NMDA receptor blockers. We bath-applied 20 μM of D-AP5, and after obtaining a baseline of 5 min, applied the standard pre + post HFS_100APs_ at 100 Hz (Fig. 3a–c). Combined HFS of the mossy fiber bouton (MFB) and the postsynaptic CA3 cell in the presence of D-AP5 did not recover PTP. EPSC_1_ amplitude was 205.7 ± 90.4 pA in control conditions and slightly dropped to 99.5 ± 32.9 pA post HFS_100_ (P = 0.13; 5 pairs; Fig. 3a–c). We therefore tested alternative Ca^2+^ sources. Application of 10 μM of the L-type Ca^2+^ channel blocker nimodipine only slightly increased the amount of PTP induced by pre + post HFS_100_ (EPSC_1_: 343.8 ± 87.4 pA in control versus 482.9 ± 161.1 pA post HFS_100_; P = 0.34; 6 pairs; Fig. 3d–f). Thus, L-type Ca^2+^ channels contributed only slightly to the relevant postsynaptic Ca^2+^ inflow. Next, we examined the possible contribution of T-type and R-type Ca^2+^ channels by applying Ni^2+^ at 100 μM (Randall et al., 1998). Applying a paired HFS_100_ at both the presynaptic terminal and the postsynaptic CA3 pyramidal cell led to a recovery of PTP (165.25 ± 85.6 pA in control versus 393.18 ± 149.5 pA post HFS_100_; P = 0.04; 7 pairs; Fig. 3g–i). Consistent with previous results (Breustedt et al., 2003; Li et al., 2007), 100 μM Ni^2+^ did not affect basal synaptic transmission. Finally, we tested the combined effects of 10 μM nimodipine and 100 μM Ni^2+^. Applying a paired HFS_100_ in the presence of both Ca^2+^ channel blockers led to a full recovery of PTP (309.2 ± 142.7 pA in control versus 988.2 ± 81.7 pA post HFS_100_; P = 0.04; 5 pairs; Fig. 3j–l). Given that T-type channels suffer from voltage-dependent inactivation around the resting potential of CA3 pyramidal neurons (−65 mV; Kim et al., 2012), our results are most consistent with the hypothesis that postsynaptic R-type Ca^2+^ channels are primarily responsible for anti-associativity triggered by combined MFB–CA3 pyramidal cell firing.

**Fig. 3.**
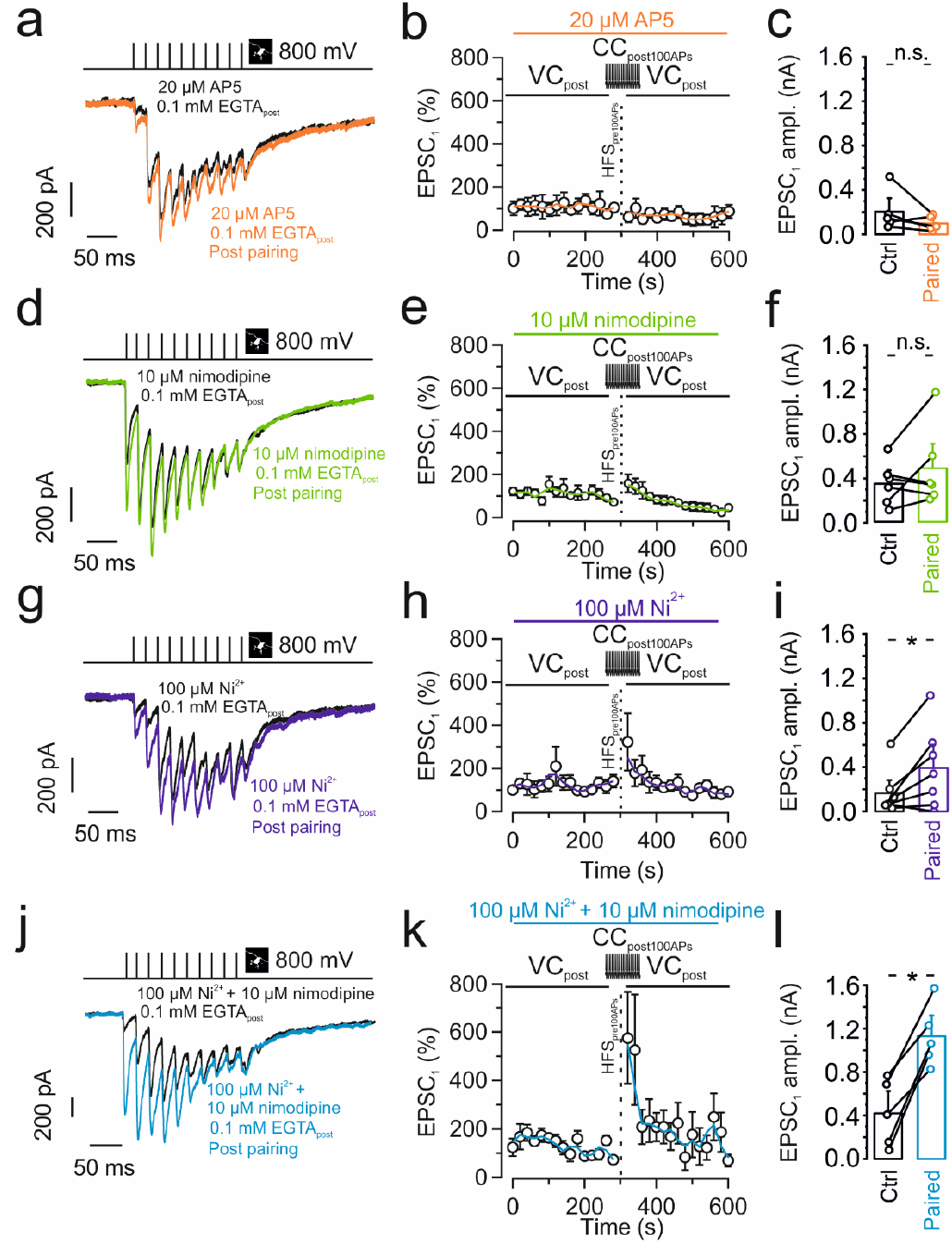
Anti-associative PTP involves postsynaptic Ca^2+^ inflow deriving from Rand L-type Ca^2+^ channels. **a–c,** Pre + post HFS_100_ in the presence of 20 μM D-AP5 does not restore PTP. **a,** representative traces of control (black) and post HFS_100_ pairing (orange). **b,** Normalized EPSC_1_ amplitude plotted against experimental time. Black vertical dashed line indicates delivery of an HFS_100_ stimulation (1 s, 100 Hz) in tight-seal cell-attached mode at the level of the MFB, combined with pairing 100 postsynaptic APs in currentclamp mode. Orange line shows the running average. Orange horizontal line on top indicates the application period of the NMDA receptor blocker D-AP5. **c,** Summary bar graph showing the EPSC_1_ amplitude before (“Ctrl”; black) and after pairing 100 presynaptic APs with postsynaptic APs (“Paired”; orange) in the presence of 20 μM D-AP5. **d–f,** Similar experiments as shown in **a–c**, but with 10 μM of the L-type Ca^2+^ channel blocker nimodipine (green). Nimodipine partially restored PTP. **g–i,** Similar experiments as shown in **a–c**, but with 100 μM of the T-type and R-type Ca^2+^ channel blocker Ni^2+^ (purple). Ni^2+^ partially restored PTP. **j–l,** Similar experiments as shown in **a–c**, but with 10 μM of the L-type Ca^2+^ channel blocker nimodipine and 100 μM of the T- or R-type Ca^2+^ channel blocker Ni^2+^ (blue). Combined application of the Ca^2+^ channel blockers completely restored PTP. **c,f,i,l,** Boxes represent mean ± SEM, a paired non-parametric Wilcoxon signed rank test was used to test for statistical significance. * indicates P < 0.05, and n.s. denotes non-significant difference (P ≥ 0.05).

Finally, we examined the molecular targets at the presynaptic terminal responsible for the apparently transsynaptic regulation of PTP (Fig. 4). At immature MFB–CA3 pyramidal neuron synapses, postsynaptic spiking is known to lead to endocannabinoid release and CB1 receptor activation at the presynaptic terminal (Caiati et al., 2012). This results in suppression of neurotransmitter release and could in theory explain our findings. However, 5 μM of the endocannabinoid receptor 1 (CB1) blocker AM251 did not significantly change the magnitude of PTP evoked by pre + post HFS_100_ (303.4 ± 130.8 pA in control versus 422.8 ± 217.9 pA post HFS_100_; P = 0.17; 6 pairs; Fig. 4a–c). Another possible mechanism would be glutamate-mediated inhibition via the presynaptic mGluR2 receptor, which is highly enriched in hippocampal mossy fiber axons and terminals (Kamiya et al., 1996; Yokoi et al., 1996; Shigemoto et al., 1997). Notably, application of 3 μM of the group II (mGluR2 / mGluR3) antagonist LY341495 partially rescued PTP evoked by pre + post HFS_100_ (273.9 ± 75.9 pA in control versus 805.1 ± 239.3 pA post HFS_100_; P = 0.005; 10 pairs; Fig. 4d–f).

**Fig. 4.**
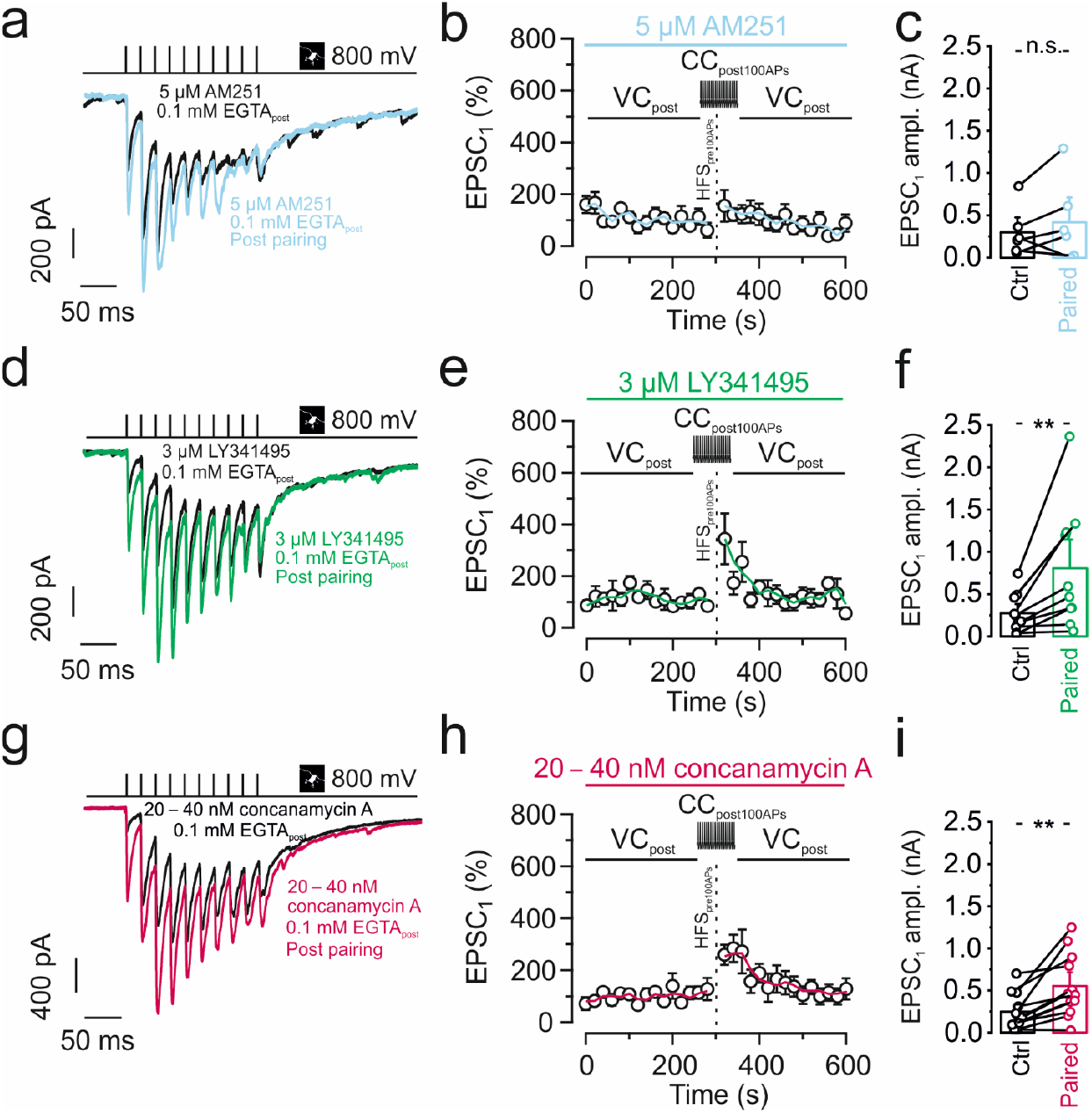
Anti-associative PTP involves retrograde glutamate signaling and activation of presynaptic mGluR2 receptors. **a–c,** Pre + post HFS_100_ in the presence of 5 μM of the CB1 receptor anatagonist AM251 does not restore PTP. **a,** Representative traces of control (black) and post HFS 100 pairing (light blue). **b,** Normalized EPSC_1_ amplitude plotted versus experimental time. Black vertical dashed line indicates delivery of an HFS_100_ stimulation (1 s, 100 Hz) in tight-seal cell-attached mode at the level of the MFB, combined with pairing 100 postsynaptic APs in current-clamp mode. Light blue line shows the running average. Light blue horizontal line on top indicates the application period of the CB1 receptor antagonist. **c,** Summary bar graph showing the EPSC_1_ amplitude before (“Ctrl”; black) and after pairing 100 presynaptic APs with postsynaptic APs (“Paired”; light blue) in the presence of 5 μM AM251. **d–f,** Pre + post HFS_100_ in the presence of 3 μM of the mGluR2 / mGluR3 antagonist LY341495 largely restored PTP. **d,** Representative traces of control (black) and post HFS 100 pairing (green). **e**, Normalized EPSC_1_ amplitude plotted versus experimental time. Black vertical dashed line indicates delivery of an HFS_100_ stimulation (1 s, 100 Hz) in tight-seal cell-attached mode at the level of the MFB, combined with pairing 100 postsynaptic APs in current-clamp mode. Green line shows the running average. Green horizontal line on top indicates the application period of the mGluR2 receptor antagonist. **f,** Summary bar graph showing the EPSC_1_ amplitude before (“Ctrl”; black) and after pairing 100 presynaptic APs with postsynaptic APs (“Paired”; green) in the presence of 3 μM LY341495. **g–i,** Pre + post HFS_100_ in the presence of 20–40 nM of the H^+^-ATPase inhibitor concanamycin A (also called folimycin) largely restores PTP. **g,** Representative traces of control (black) and post HFS_100_ pairing (dark green). **h**, Normalized EPSC_1_ amplitude plotted versus experimental time. Black vertical dashed line indicates delivery of an HFS_100_ stimulation (1 s, 100 Hz) in tight-seal cell-attached mode at the level of the MFB, combined with pairing 100 postsynaptic APs in current-clamp mode, magenta line shows the running average. Magenta horizontal line on top indicates the application period of the H^+^-ATPase inhibitor. **f,**Summary bar graphs showing the EPSC_1_ amplitude before (“Ctrl”; black) and after pairing 100 presynaptic APs with postsynaptic APs (“Paired”; magenta) in the presence of 20–40 nM of concanamycin A. **c,f,i** Boxes represent mean ± SEM, a paired non-parametric Wilcoxon signed rank test was used to test for statistical significance. * indicates P < 0.05, and n.s. denotes nonsignificant difference (P ≥ 0.05).

Given that the anti-associative mechanism was dependent on postsynaptic Ca^2+^ chelators, we tested the contribution of retrograde vesicular glutamate signaling. Since glutamate transport into vesicles typically requires a proton gradient, we reasoned that interfering with this mechanism might mimic the effect of the mGluR2 / mGluR3 antagonist LY341495. To specifically target the postsynaptic neuron, we loaded CA3 pyramidal neurons with the v-type H^+^-ATPase inhibitor concanamycin A (20–40 nM) via the patch pipette. Postsynaptic specificity is demonstrated by the lack of inhibition of presynaptic mediated neurotransmitter release (baseline, Fig. 4h). Interestingly, treatment with concanamycin A led to a moderate but highly significant recovery of PTP (246.8 ± 62.5 pA in control versus 553.9 ± 112.3 pA post HFS_100_; P = 0.003; 12 pairs; Fig. 4g–i). Taken together, our results may suggest a retrograde signaling mechanism, in which postsynaptic activity induces an increase in postsynaptic Ca^2+^ concentration, vesicular release of a retrograde messenger, probably glutamate, from the dendrites of the postsynaptic cell (Glitsch et al., 1996; Zilberter, 2000), and subsequent activation of presynaptic mGluRs (Fig. 5).

**Fig. 5.**
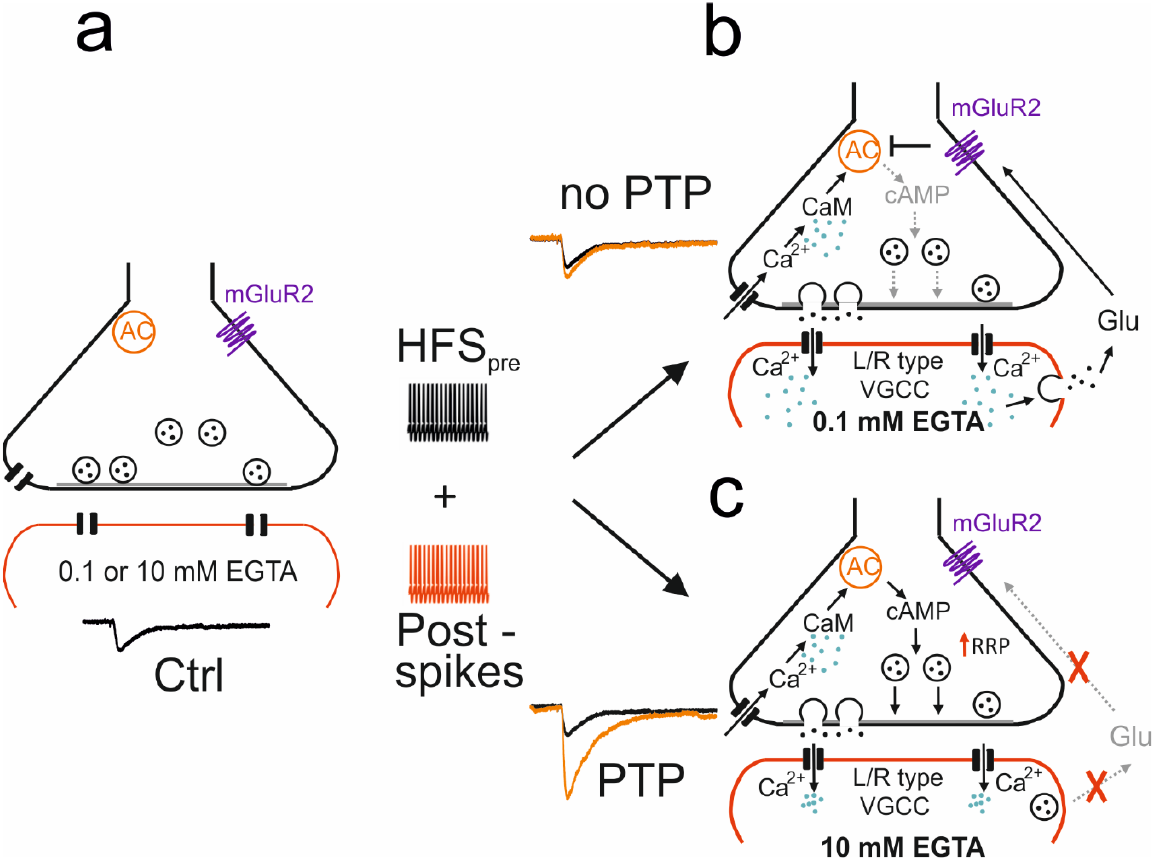
Schematic summary of the induction rules and underlying mechanisms of PTP at the MFB–CA3 pyramidal neuron synapse. **a,** MFB–CA3 synapse during resting conditions. Presynaptic terminal in black on top, postsynaptic compartment in red at the bottom. AC stands for adenylyl cyclase, mGluR2 indicates extrasynaptic metabotropic glutamate receptor 2 (and mGluR3 expressed at lower levels). **b,** MFB–CA3 synapse after tetanic stimulation, leading to post-synaptic activity, in the presence of low (0.1 mM) concentrations of the Ca^2+^ chelator EGTA. CaM stands for calmodulin, VGCC stands for voltage-gated Ca^2+^ channels, and Glu stands for glutamate. Ca^2+^ ions indicated as green dots, glutamate molecules as black dots. Note that in this scenario tetanic stimulation does not trigger PTP, due to retrograde vesicular glutamate signaling. **c,** Same as for b but in the presence of high concentration (10 mM) of the Ca^2+^ chelator EGTA. Note that in this scenario tetanic stimulation does trigger PTP, due to the lack of retrograde vesicular glutamate signaling. Transsynaptic modulation of PTP induction may not only occur with the exogenous Ca^2+^ chelator EGTA, but also with endogenous Ca^2+^ buffers expressed in the postsynaptic neurons. This may convey both anti-associativity and target cell-specificity to the PTP induction mechanism (Alle et al., 2001).

## DISCUSSION

Synapse specificity, cooperativity, and associativity are hallmark features of synaptic plasticity at glutamatergic synapses. However, rigorous testing of these properties is often difficult, because it requires defined stimulation of individual synaptic inputs. To determine specificity, cooperativity, and associativity of hippocampal mossy fiber PTP, we combined paired recording between mossy fiber terminals and postsynaptic CA3 pyramidal neurons in rat brain slices (Vyleta et al., 2016; Vandael et al., 2020) with extracellular tract stimulation. This allowed us to stimulate single inputs, as required for cooperativity analysis, to validate the non-overlapping nature of the inputs by presynaptic action current measurements, essential for specificity experiments, and to precisely control pre- and postsynaptic activity, critically important for characterization of associativity.

Using this tightly controlled approach, we demonstrate that hippocampal mossy fiber PTP shows synapse specificity, but lacks cooperativity and associativity. PTP induction is not only non-associative, but rather shows anti-associative properties. Non-cooperativity might have been expected, given that PTP can be robustly induced by single bouton stimulation (Vyleta et al., 2016; Vandael et al., 2020). In contrast, the anti-associativity was surprising. Although previous work showed that mossy fiber synaptic plasticity can be enhanced by postsynaptic hyperpolarization (Sokolov et al., 2003), this is, to the best of our knowledge, the first description of anti-associative properties of short-term plasticity at a glutamatergic synapse. The unique induction rules of PTP provide the synapse with complete autonomy and a powerful feedback mechanism to prevent excessive detonation in the hippocampal network.

Our results show that anti-associativity can be observed under conditions in which postsynaptic HFS_100_ is induced by postsynaptic current injections and, albeit to a smaller extent, conditions in which postsynaptic CA3 pyramidal neuron freely fires in response to mossy fiber terminal simulation during PTP induction. This corroborates the physiological significance of the observed anti-associativity mechanism. Interestingly, the suppressive effect on PTP is better correlated with the coefficient of variation of the spike interval than with the number of APs. This suggests that the anti-associativity requires, or is facilitated by, burst firing in the postsynaptic neuron. This is consistent with the involvement of R-type Ca^2+^ channels, which has been reported to promote burst firing in subicular neurons (Metz et al., 2005, see below).

What are the mechanisms underlying anti-associative PTP induction? Our results show that both Ca^2+^ chelators introduced into the postsynaptic cell and L- and R-type Ca^2+^ channel blockers, particularly when applied in combination, rescued PTP. Thus, Ca^2+^ inflow through postsynaptic voltage-gated Ca^2+^ channels plays a critical role in the suppression of PTP. This is consistent with the presence of high-voltage activated, Ni^2+^-sensitive Ca^2+^ channels (putatively R-type) in the dendrites of hippocampal pyramidal neurons (Magee and Johnston, 1995). Earlier work demonstrated that mossy fiber PTP is generated by an increase in the size of the readily releasable synaptic vesicle pool (Vandael et al., 2020). As these mechanisms operate in the presynaptic terminals, retrograde transsynaptic signaling mechanisms must be involved (Regehr et al., 2009). Our results identify mGluR2 or mGluR3, probably activated by dendritically released glutamate, as the critical link between postsynaptic spiking activity and presynaptic terminal function. mGluR2 and mGluR3 are known to couple to Gi and thereby inhibit adenylyl cyclase (Kamiya et al., 1996; Conn and Pin, 1997). Adenylyl cyclase, in turn, regulates the size of the readily releasable pool in mossy fiber terminals (Vandael et al., 2020). Thus, postsynaptic activity could regulate presynaptic vesicle pool size (Fig. 5).

How exactly glutamate is released from postsynaptic CA3 cells remains unclear. Our finding that the vacuolar-type H^+^-ATPase inhibitor concanamycin A introduced in the postsynaptic pipette reduced the suppression of PTP may be consistent with a vesicular release mechanism, in which glutamate accumulation depends on established proton gradients. Consistent with this hypothesis, thorny excrescences of CA3 pyramidal neurons show a pronounced spine apparatus and are enriched in multivesicular bodies (Chicurel and Harris, 1992). More work is needed to disentangle the role of these structures in retrograde signaling. If dendritically released glutamate inhibits PTP via mGluR2 or 3, why does glutamate released from presynaptic terminals not have the same effect? MGluR2 or 3 show a relatively low affinity for glutamate (Conn and Pin, 1997). Furthermore, mGluR2 is primarily located in extrasynaptic regions of mossy fiber axons (Shigemoto et al., 1997). In combination, receptor properties and subcellular localization may facilitate activation of the pathway by dendritic release.

Previous work identified several mechanisms that enhance synaptic efficacy at hippocampal mossy fiber synapses, including facilitation, post-tetanic potentiation, and long-term potentiation (Salin et al., 1996; Toth et al., 2000; Vyleta and Jonas, 2014; Vyleta et al., 2016). This led to the suggestion that the hippocampal mossy fiber– CA3 pyramidal neuron synapse can operate as a conditional or plasticity-dependent full detonator (Treves and Rolls, 1992; Henze et al., 2002; Pelkey and McBain, 2005; Vyleta et al., 2016). However, the abundance of multiple forms of potentiating plasticity raises the question of how these detonation mechanisms might be controlled. Transsynaptic modulation of PTP induction may represent a simple, but powerful mechanism to prevent excessive potentiation in the hippocampal network, and to implement a break on mossy fiber detonation. This may be important to keep the balance between excitation and inhibition in the network.

The present findings may have major implications for the memory function of the hippocampal network. A prevalent model of hippocampal memory suggests that mossy fiber synapse acts as a “teacher” that triggers the storage and recall of memories in the downstream CA3 region (Rolls, 2016). In this model, the mossy fiber synapse is often viewed as a conditional detonator, in which the efficacy of the synapse depends on prior presynaptic activity (Henze et al., 2002; Vyleta et al., 2016). Our results challenge this view, showing that the detonation properties of the synapse depend on both pre- and postsynaptic activity in a complex manner. This might be a powerful mechanism to ensure that storage and recall are separated and that new information is preferentially stored in silent, non-coding CA3 pyramidal neurons (Kowalski et al., 2016). Thus, the mossy fiber synapse may act as a “smart teacher”, which may help to maximize the storage capacity in the system. Future experiments combining electrophysiological analysis, network modeling, and behavioral analysis will be needed to further test this hypothesis.

## ACKNOWLEDGMENTS

We thank Drs. Carolina Borges-Merjane and Jose Guzman for critically reading the manuscript, and Pablo Castillo for useful discussions. We are grateful to Alois Schlögl for help with analysis, Florian Marr for excellent technical assistance and cell reconstruction, Christina Altmutter for technical help, Eleftheria Kralli-Beller for manuscript editing, and the Scientific Service Units of IST Austria for support. This project received funding from the European Research Council (ERC) under the European Union’s Horizon 2020 research and innovation program (grant agreement No 692692) and the Fond zur Förderung der Wissenschaftlichen Forschung (Z 312-B27, Wittgenstein award), both to P.J.

## AUTHOR CONTRIBUTIONS

D.V. and Y.O. performed the experiments and analyzed the data, P.J. and D.V. conceived the project and wrote the paper. All authors analyzed data and jointly revised the paper.

## Supplementary Methods

### Animal experiments

Paired pre- and postsynaptic recordings in vitro were carried out on 17- to 23-day-old Wistar rats (RRID:RGD_13508588; weight: 55–65 g). Animals were housed under a reversed light cycle (dark: 7:00 am – 7:00 pm, light: 7:00 pm – 7:00 am). For experiments, both male and female animals were used. All experiments were carried out in strict accordance with institutional, national, and European guidelines for animal experimentation, and approved by the Bundesministerium für Wissenschaft, Forschung und Wirtschaft of Austria (A. Haslinger).

### Paired recordings from mossy fiber terminals

Transverse hippocampal slices (350–400 μm thick) were prepared according to previously published protocols (Bischofberger et al., 2006; Vandael et al., 2020). Animals were anesthetized using isoflurane and killed by rapid decapitation. Slices were cut from the right or left hemisphere in ice-cold, sucrose-containing extracellular solution using a vibratome (VT1200, Leica Microsystems), incubated in a maintenance chamber at ~35°C for 30–45 min, and subsequently stored at room temperature. Cutting solution contained 64 mM NaCl, 25 mM NaHCO_3_, 2.5 mM KCl, 1.25 mM NaH_2_PO_4_, 10 mM glucose, 120 mM sucrose, 0.5 mM CaCl_2_, and 7 mM MgCl_2_. Storage solution contained 87 mM NaCl, 25 mM NaHCO_3_, 2.5 mM KCl, 1.25 mM NaH_2_PO_4_, 10 mM glucose, 75 mM sucrose, 0.5 mM CaCl_2_, and 7 mM MgCl_2_ (equilibrated with 95% O_2_ and 5% CO_2_). Experiments were performed at room temperature (24.1 ± 0.2 °C; range: 21–26°C. Before onset of the experiment, slices were placed in the recording chamber and superfused with artificial cerebrospinal fluid (ACSF; 125 mM NaCl, 25 mM NaHCO_3_,, 2.5 mM KCl, 1.25 mM NaH_2_PO_4_, 2 mM CaCl_2_, and 1 mM MgCl_2_, equilibrated with 95% O_2_ and 5% CO_2_) for at least 15 minutes before recording.

Subcellular patch-clamp recordings from MFBs and simultaneous recordings from pyramidal neurons in the CA3b and CA3c region of the hippocampus were performed under visual control provided by infrared differential interference contrast videomicroscopy (Bischofberger et al., 2006; Vandael et al., 2020). Presynaptic and postsynaptic recording pipettes were fabricated from borosilicate glass tubing (2.0 mm outer diameter, 1.0 mm inner diameter) and had open-tip resistances of 10–20 MΩ and 3–7 MΩ, respectively. For tight-seal, bouton-attached stimulation under voltage-clamp conditions, the presynaptic pipette contained a K^+^-based intracellular solution (130 mM K-gluconate, 2 mM KCl, 2 mM MgCl_2_, 2 mM Na_2_ATP, 10 mM HEPES, 10 mM EGTA, and 0.2% biocytin; or 140 mM KCl, 2 mM MgCl_2_, 4 mM Na_2_ATP, 0.3 mM Na_2_GTP, 10 mM HEPES, 0.1 mM EGTA, and 0.2% biocytin; pH adjusted to 7.28 with KOH), allowing us to minimally invasively stimulate boutons in the tight-seal cell-attached configuration. In the cell-attached configuration, seal resistance was > 1 GΩ and holding potential was set at –70 mV to minimize the holding current. APs in MFBs were evoked by brief voltage pulses (amplitude 800 mV, duration 0.1 ms). Mossy fiber terminals had diameters of ~2–5 μm, in agreement with the previously reported range of diameters of MFBs in light and electron microscopy studies. For PTP time course experiments, stimuli (10 stimuli at 50 Hz) were delivered once every 20 s. For tract stimulation experiments, a large pipette (tip diameter 5–10 μm) filled with 1 M NaCl was placed in the subgranular zone of the dentate gyrus. Stimulation was performed using a stimulus isolation unit; simulation intensity was ~30 V (range: 10–100 V).

Postsynaptic recording pipettes contained an internal solution containing 130 mM K-gluconate, 2 mM or 20 mM KCl, 2 mM MgCl_2_, 2 mM Na_2_ATP, 10 mM HEPES, 10 mM or 0.1 mM EGTA, and 0.2% biocytin (pH adjusted to 7.28 with KOH). In a subset of recordings 0.3 mM GTP was added to a postsynaptic internal solution, and 20 mM glucose was added to adjust the osmolarity in K^+^-based intracellular solution with 2 mM KCl. For whole-cell voltage-clamp recordings from postsynaptic CA3 pyramidal neurons, the membrane potential was set to −70 or −80 mV. Only recordings with < 300 pA leakage current were included in the analyses. Postsynaptic series resistance was kept below 10 MΩ (8.25 ± 0.26 MΩ, 116 mossy fiber terminal– CA3 pyramidal neuron recordings). In the voltage-clamp mode, series resistance was uncompensated, but carefully monitored with a test pulse following each data acquisition sweep. Only recordings with stable series resistance (with < 20% change during the recording period) were included in the analyses. Paired recordings (presynaptic tight-seal, bouton-attached stimulation, postsynaptic whole-cell recording) were stable for up to 30 min. Nimodipine, D-AP5, AM251, and concanamycin A were dissolved in dimethyl sulfoxide (DMSO) at concentrations of 10 mM, 10 mM, 10 mM, and 20 μM, respectively, and then added to the ACSF. LY341495 (free base) was dissolved in DMSO, LY341495 (disodium salt) was dissolved in water at a concentration of 6 mM. Free base and disodium salt of LY341495 gave identical results, therefore data were pooled. The final concentration of DMSO in the ACSF was ≤ 0.2%. Nimodipine, D-AP5, AM251, concanamycin A, and LY341495 were from Tocris, other chemicals were from Sigma-Aldrich.

### Data analysis

Data were acquired with a Multiclamp 700A amplifier, low-pass filtered at 10 kHz, and digitized at 40- or 100 kHz using a CED power1401 plus or mkII interface (Cambridge Electronic Design, Cambridge, UK). Pulse generation and data acquisition were performed using FPulse version 3.3.3 (U. Fröbe, Physiological Institute, University of Freiburg, Germany). Data were analyzed using Stimfit version 0.14 or 0.15 (Guzman et al., 2014) and Igor Pro 6.37 (Wavemetrics). For peak detection, the analysis time window was set between 1 and 19 ms after the stimulus. EPSC decay time course was fit with a mono-exponential function plus an offset. Similarly, PTP decay time course was fit with a mono-exponential function plus an offset. Since PTP peaked in between 20 s and 60 s after HFS_100_, we compared the peak amplitude of the largest EPSC in this time interval with that of the average EPSC in a time interval 100 s before HFS, which was defined as PTP_max_. For Fig. 2h and 2i, we used the average of PTP_max_ of the first three EPSCs to enhance the reliability of recordings.

### Statistics and conventions

Statistical significance was assessed using a two-sided Wilcoxon signed rank test for paired comparisons or a Mann-Whitney U test for unpaired comparisons at the significance level (*P*) indicated, using Python 2. Multiple comparisons were performed with a Kruskal-Wallis test. Correlation analysis was performed with Spearman’s rank test. Values are given as mean ±standard error of the mean (SEM). Error bars in the Figures also represent the SEM. For graphical representation of statistics, * indicates P < 0.05, ** P < 0.01, and *** P < 0.001. Membrane potentials are given without correction for liquid junction potentials. In total, data reported in this paper were obtained from 116 paired mossy fiber terminal–CA3 pyramidal neuron recordings. 12 of these recordings (Fig. 2g, VCnoAPs) were already reported in a previous publication (Vandael et al., 2020).

### Data availability

Data from this study are available from the corresponding authors upon reasonable request.

### Code availability

Analysis routines and code are available from the corresponding authors upon reasonable request.

## References

1. Abbott, L.F. & Regehr, W.G. Synaptic computation. Nature 431, 796–803 (2004).

2. Alle, H., Jonas, P. & Geiger, J.R.P. PTP and LTP at a hippocampal mossy fiber interneuron synapse. Proc. Natl. Acad. Sci. USA 98, 14708–14713 (2001).

3. Bischofberger, J., Engel, D., Li, L., Geiger, J.R.P. & Jonas, P. Patch-clamp recording from mossy fiber terminals in hippocampal slices. Nat. Protoc. 1, 2075–2081 (2006).

4. Breustedt, J., Vogt, K.E., Miller, R.J., Nicoll, R.A. & Schmitz, D. Alpha1E-containing Ca^2+^ channels are involved in synaptic plasticity. Proc. Natl. Acad. Sci. USA 100, 12450–12455 (2003).

5. Brown, T.H. & Johnston, D. Voltage-clamp analysis of mossy fiber synaptic input to hippocampal neurons. J. Neurophysiol. 50, 487–507 (1983).

6. Caiati, M.D., Sivakumaran, S, Lanore, F., Mulle, C., Richard, E., Verrier, D., Marsicano, G., Miles, R. & Cherubini E. Developmental regulation of CB1-mediated spike-time dependent depression at immature mossy fiber–CA3 synapses. Sci. Rep. 2, 285 (2012).

7. Chicurel, M.E. & Harris, K.M. Three-dimensional analysis of the structure and composition of CA3 branched dendritic spines and their synaptic relationships with mossy fiber boutons in the rat hippocampus. J. Comp. Neurol. 325, 169–182 (1992).

8. Conn, P.J. & Pin, J.P. Pharmacology and functions of metabotropic glutamate receptors. Annu. Rev. Pharmacol. Toxicol. 37, 205–237 (1997).

9. Erickson, M.A., Maramara, L.A. & Lisman, J. A single brief burst induces GluR1-dependent associative short-term potentiation: a potential mechanism for shortterm memory. J. Cogn. Neurosci. 22, 2530–2540 (2010).

10. Fiebig, F. & Lansner, A. A spiking working memory model based on Hebbian short-term potentiation. J. Neurosci. 37, 83–96 (2017).

11. Glitsch, M., Llano, I. & Marty, A. Glutamate as a candidate retrograde messenger at interneurone-Purkinje cell synapses of rat cerebellum. J. Physiol. 497, 531 – 537 (1996).

12. Griffith, W.H. Voltage-clamp analysis of posttetanic potentiation of the mossy fiber to CA3 synapse in hippocampus. J. Neurophysiol. 63, 491–501 (1990).

13. Guzman, S.J., Schlögl, A. & Schmidt-Hieber, C. Stimfit: quantifying electrophysiological data with Python. Front. Neuroinform. 8, 16 (2014).

14. Henze, D.A., Urban, N.N. & Barrionuevo, G. The multifarious hippocampal mossy fiber pathway: a review. Neuroscience 98, 407–427 (2000).

15. Henze, D.A., Wittner, L. & Buzsáki, G. Single granule cells reliably discharge targets in the hippocampal CA3 network *in vivo*. Nat. Neurosci. 5, 790–795 (2002).

16. Jonas, P., Major, G. & Sakmann, B. Quantal components of unitary EPSCs at the mossy fibre synapse on CA3 pyramidal cells of rat hippocampus. J. Physiol. 472, 615–663 (1993).

17. Kamiya, H., Shinozaki, H. & Yamamoto C. Activation of metabotropic glutamate receptor type 2/3 suppresses transmission at rat hippocampal mossy fibre synapses. J. Physiol. 493, 447–455 (1996).

18. Kwon HB, Castillo PE. Long-term potentiation selectively expressed by NMDA receptors at hippocampal mossy fiber synapses. Neuron 57, 108–120 (2008).

19. Kim, S., Guzman, S.J., Hu, H. & Jonas, P. Active dendrites support efficient initiation of dendritic spikes in hippocampal CA3 pyramidal neurons. Nat. Neurosci. 15, 600–606 (2012).

20. Kowalski, J., Gan, J., Jonas, P. & Pernía-Andrade, A.J. Intrinsic membrane properties determine hippocampal differential firing pattern in vivo in anesthetized rats. Hippocampus 26, 668–682 (2016).

21. Langdon, R.B., Johnson, J.W. & Barrionuevo, G. Posttetanic potentiation and presynaptically induced long-term potentiation at the mossy fiber synapse in rat hippocampus. J. Neurobiol. 26, 370–385 (1995).

22. Li, L., Bischofberger, J. & Jonas, P. Differential gating and recruitment of P/Q-, N-, and R-type Ca^2+^ channels in hippocampal mossy fiber boutons. J. Neurosci. 27, 13420–13429 (2007).

23. Magee, J.C. & Johnston, D. Characterization of single voltage-gated Na^+^ and Ca^2+^ channels in apical dendrites of rat CA1 pyramidal neurons. J. Physiol. 487, 67–90 (1995).

24. Metz, A.E., Jarsky, T., Martina, M. & Spruston, N. R-type calcium channels contribute to afterdepolarization and bursting in hippocampal CA1 pyramidal neurons. J. Neurosci. 25, 5763–5773.

25. Mishra, R.K., Kim, S., Guzman, S.J. & Jonas, P. Symmetric spike timingdependent plasticity at CA3–CA3 synapses optimizes storage and recall in autoassociative networks. Nat. Commun. 7, 11552 (2016).

26. Nicoll, R.A. & Schmitz, D. Synaptic plasticity at hippocampal mossy fibre synapses. Nat. Rev. Neurosci. 6, 863–876 (2005).

27. Pelkey, K.A. & McBain C.J. How to dismantle a detonator synapse. Neuron 45, 327–329 (2005).

28. Randall, A.D. The molecular basis of voltage-gated Ca^2+^ channel diversity: is it time for T? J. Membr. Biol. 161, 207–213 (1998).

29. Regehr, W.G., Carey, M.R. & Best, A.R. Activity-dependent regulation of synapses by retrograde messengers. Neuron 63, 154–170 (2000).

30. Rolls, E.T. Pattern separation, completion, and categorisation in the hippocampus and neocortex. Neurobiol. Learn. Mem. 129, 4–28 (2016).

31. Salin, P.A., Scanziani, M., Malenka, R.C. & Nicoll, R.A. Distinct short-term plasticity at two excitatory synapses in the hippocampus. Proc. Natl. Acad. Sci. USA 93, 13304–13309 (1996).

32. Sastry, B.R., Goh, J.W. & Auyeung, A. Associative induction of posttetanic and long-term potentiation in CA1 neurons of rat hippocampus. Science 232, 988–990 (1986).

33. Schmitz, D., Mellor, J., Breustedt, J. & Nicoll R.A. Presynaptic kainate receptors impart an associative property to hippocampal mossy fiber long-term potentiation. Nat. Neurosci. 6, 1058–1063 (2003).

34. Shigemoto, R., Kinoshita, A., Wada, E., Nomura, S., Ohishi, H., Takada, M., Flor, P.J., Neki, A., Abe, T., Nakanishi, S. & Mizuno, N. Differential presynaptic localization of metabotropic glutamate receptor subtypes in the rat hippocampus. J. Neurosci. 17, 7503–7522 (1997).

35. Silver, R.A. Neuronal arithmetic. Nat. Rev. Neurosci. 11, 474–489 (2010).

36. Sokolov, M.V., Rossokhin, A.V., M. Kasyanov, A., Gasparini, S., Berretta, N., Cherubini, E. & Voronin, L.L. Associative mossy fibre LTP induced by pairing presynaptic stimulation with postsynaptic hyperpolarization of CA3 neurons in rat hippocampal slice. Eur. J. Neurosci. 17, 1425–1437 (2003).

37. Toth, K., Suares, G., Lawrence, J.J., Philips-Tansey, E. & McBain, C.J. Differential mechanisms of transmission at three types of mossy fiber synapse. J. Neurosci. 20, 8279–8289 (2000).

38. Treves, A. & Rolls, E.T. Computational constraints suggest the need for two distinct input systems to the hippocampal CA3 network. Hippocampus 2, 189–199 (1992).

39. Vandael, D., Borges-Merjane, C., Zhang, X. & Jonas, P. Short-term plasticity at hippocampal mossy fiber synapses is induced by natural activity patterns and associated with vesicle pool engram formation. Neuron 107, 509–521.e7 (2020).

40. Vyleta, N.P. & Jonas, P. Loose coupling between Ca^2+^ channels and release sensors at a plastic hippocampal synapse. Science 343, 665–670 (2014).

41. Vyleta, N.P., Borges-Merjane, C. & Jonas, P. Plasticity-dependent, full detonation at hippocampal mossy fiber–CA3 pyramidal neuron synapses. eLife 5, e17977 (2016).

42. Weisskopf, M.G. & Nicoll, R.A. Presynaptic changes during mossy fibre LTP revealed by NMDA receptor-mediated synaptic responses. Nature 376, 256–259 (1995).

43. Yokoi, M., Kobayashi, K., Manabe, T., Takahashi, T., Sakaguchi, I., Katsuura, G., Shigemoto, R., Ohishi, H., Nomura, S., Nakamura, K., Nakao, K., Katsuki, M. & Nakanishi, S. Impairment of hippocampal mossy fiber LTD in mice lacking mGluR2. Science 273, 645–647 (1996).

44. Zalutsky, R.A. & Nicoll, R.A. Mossy fiber long-term potentiation shows specificity but no apparent cooperativity. Neurosci. Lett. 138, 193–197 (1992).

45. Zilberter, Y. Dendritic release of glutamate suppresses synaptic inhibition of pyramidal neurons in rat neocortex. J. Physiol. 528, 489–496 (2000).

